# Impact of Zinc Pre-exposure on de novo Antibiotic Resistance Development

**DOI:** 10.1101/2023.04.10.536219

**Authors:** Mark P. Suprenant, Carly Ching, Indorica Sutradhar, Neila Gross, Jessica E. Anderson, Nourhan El Sherif, Muhammad H. Zaman

**Affiliations:** Department of Biomedical Engineering, Boston University, Boston Massachusetts, USA; Department of material Science Engineering, Boston University, Boston Massachusetts, USA; Howard Hughes Medical Institute, Boston University, Boston Massachusetts, USA; Center on Forced Displacement, Boston University, Boston Massachusetts, USA

**Author notes:** Address correspondence to Muhammad Zaman.

## Abstract

Antimicrobial resistance (AMR) is a global health crisis that is currently predicted to worsen. While the impact of improper antibiotics is an established driver, much less is known on the impacts of metal supplements. Here, we specifically probe the impact of zinc (Zn) on AMR. In conflict settings where diarrhea disease cases are high, Zn is both given as a supplement for treatment of these diseases prior to use of antibiotics such as ciprofloxacin and is associated with weapons of war. In this study, we find that the order with which *E. coli* is exposed to zinc impacts resistance development, with increasing pre-exposure time leading to accelerated ciprofloxacin resistance, while combined exposure of zinc with ciprofloxacin delays ciprofloxacin resistance. We did not find evidence that zinc pre-exposure leads to genetic changes or change in antibiotic tolerance, though it does increase both the lag phase and doubling time of *E. coli*, suggesting the mechanism may be due to changes in gene expression. While the zinc phenotype behavior is not permeant and would no longer be observed if ciprofloxacin exposure did not occur right after zinc pre-exposure, the elevated MIC phenotype resulting from the zinc pre-exposure was more stable than the zinc phenotype. These results are important as they highlight the need to reexamine the clinical role of zinc in treating diarrheal diseases and assess if changes in resistance development observed in vitro are also observed in vivo.

**Importance:** Antimicrobial resistance (AMR) is a global problem. According to a 2014 Review on Antimicrobial Resistance, it is projected to result in several million deaths by 2050 (Review on Antimicrobial Resistance, Tackling a Crisis for the Health and Wealth of Nations, 2014). While the improper usage of antibiotic treatments is an accepted driver of AMR, little work has focused on how non-antibiotic medication, such as supplements, might impact this when combined with antibiotics. One supplement of interest is the heavy metal zinc which is used in conjunction with ciprofloxacin to treat diarrheal diseases in children. We find that the order and duration of zinc exposure has significant impact on resistance development. More specifically, although the combined presence of zinc and ciprofloxacin delays the onset of resistance, when used successively as they often are in practice, zinc pre-exposure followed by ciprofloxacin exposure results in faster resistance development.

## Introduction

The development of antimicrobial resistance (AMR) is a global health crisis. It is estimated that by 2050, AMR may account for 10 million deaths and a cost of $300 billion – $1 trillion annually^1^. The impacts of AMR are already making themselves known, however, as it is estimated that in 2019 1.27 million deaths were directly caused by AMR with an additional 3.68 million deaths associated with AMR^2^. As such, much research has been focused on investigating drivers of AMR, from the role of the veterinary sector to the impact of substandard antibiotics and sub-inhibitory antibiotic concentrations^3,4^. Beyond known pathways of drug resistance due to changes in mutation rates or horizontal gene transfer, bacterial antibiotic resistance can also develop due to exposure to other substances in the environment^5^. One such class of materials is heavy metals, which can readily contaminate aqueous environments and once they have contaminated an environment, are easily absorbed by living organisms due to their highly soluble nature^6,7^. Heavy metal contaminants traditionally arise from industrial runoff but can also be associated with weapons of war^8^. Metals like lead and mercury (which are used in explosives) as well as zinc, copper, nickel, and chromium (used to coat a range of military objects from bullets to large military vehicles), are potentially related to the development of resistance in bacterial species such as *A. baumannii*^8^.

Zinc is of particular interest since unlike other heavy metals that are needed in small doses for nutrition purposes, zinc is also given as a supplement (often in the salt form of zinc sulfate) as part of the recommended treatment protocol for diarrheal diseases in children under five years old^9^. Diarrheal diseases have a hosts of causes, though Rotavirus and *E. coli* are the most common agents of moderate to severe diarrhea in low-income countries according to the World Health Organization (https://www.who.int/news-room/fact-sheets/detail/diarrhoeal-disease). Regardless of the cause, zinc treatment is given in the form of zinc and oral rehydration salts (Zn/ORS), which at $0.50 USD per dose, is a cost-effective treatment^10^. The ORS serves to treat and prevent dehydration while the zinc has been shown to improve the response of the immune system^10–13^. Computational models have also shown a decrease in the severity and duration of diarrheal episodes despite significant reductions not being observed in earlier, small scale clinical studies^14–17^. Thus, this treatment helps improve recovery from the diarrheal disease and its associated zinc depleted state^15^, while also serving to stave off further infection and malnutrition, which based on reports by the World Health Organization could ultimately lead to stunted development, wasting, increased risk of mortality, and a decline in cognitive potential (https://www.who.int/news-room/fact-sheets/detail/malnutrition).

While the benefits of zinc supplementation for addressing diarrheal disease symptoms^16^ and risks about zinc’s impact on resistance development in the environment have been observed^18^, much less is known on how zinc supplementation treatment might impact the evolution of AMR development in bacteria found in the human gut that can cause diarrheal diseases, such as *E. coli*. Current studies largely focus on the agriculture sector, looking at the impact that zinc in feed has on animals and their gut microbiota or on the selective pressure against resistance that zinc exerts on already antibiotic resistant *E. coli*^19,20^. Literature on *de novo* resistance development is limited and presents conflicting viewpoints, such as work relating to pig feeding stating that the presence of zinc leads to larger resistant populations in some cases^21^, while in others, zinc has no impact on antibiotic reisstance^22^. Meanwhile other *in vitro* studies have reported that the presence of zinc can inhibit the development of antibiotic resistance^23^.

Understanding the interplay between zinc and bacterial resistance development is critical, especially in places where a high prevalence of diarrheal diseases, zinc supplementation, and war overlap, such as in a conflict and humanitarian emergency settings like Yemen. Given the difficulty in treating susceptible versions of these disease-causing bacteria in a conflict zone, it is imperative that current treatment practices do not further promote the development of resistant infections in these populations. Humanitarian actions should avoid perpetuating any of war’s impact on the health landscape in these countries long after the active fighting has ceased. As such, the objectives of this study were to examine the impact that zinc has on the *in vitro* development of resistance to ciprofloxacin in *E. coli*, an antibiotic frequently proscribed and taken for various diseases, including diarrheal diseases, in Yemen (Personal Communication, staff at UNICEF Yemen). More specifically we first probed if zinc sulfate has any interaction with ciprofloxacin’s ability to inhibit bacterial growth. Next, we checked to see if zinc sulfate impacted development of resistance to ciprofloxacin. Lastly, we examined how zinc sulfate exposure timing may factor into the development of resistance, as well as the stability of these impacts. Overall, our findings may provide important insights into the historical and continued practice of zinc ORS treatments.

## Methods

### Strains and culture conditions

*E. coli* MG1655 (ATCC 700926) was used in all experiments. All liquid cultures were grown in LB medium under shaking conditions at 180 rpm at 37 °C. Ciprofloxacin and zinc (Zn) in the form of zinc sulfate was added to the medium as indicated below.

### Antibiotic and Zn susceptibility testing

To determine the Minimum Inhibitory Concentration (MIC) of bacterial cells post exposure, cells were outgrown in drug-free media (not amended with Zn) until saturated (~24 hours) and subsequently used in a standard broth microdilution MIC in a 96-well plate using LB media^24^.

This assay was performed against ciprofloxacin or Zn in the form of Zn sulfate to determine the respective MICs for each substance.

To test for interactions between Zn and ciprofloxacin that impact the inhibitory concentration, a checkerboard assay was also performed as described by Bellio et. al^25^. Briefly, this assay swept over the combination of concentration values of both Zn and ciprofloxacin together over the range where the MIC for each substance would occur individually. Based on the observed inhibitory concentrations of the combined and individual drugs, we determined the fractional inhibitory concentration (FIC) of each drug to find if the drugs are synergistic (FIC>4), antagonistic (FIC<0.5) or additive/ indifferent (0.5<FIC<4)^25^.

### Zn pre-exposure time course and Zn-ciprofloxacin combination assays

At the start of the 11-day Zn exposure time experiments, saturated liquid cultures were diluted 1:100 in 4 mL of LB broth and grown for two hours to exponential phase. To start Zn preexposure, cells were then transferred at a 1:100 dilution into 4 mL of LB broth containing an additional 0.5 mM of Zn five days before the addition of antibiotics (day-5) or three days before the addition of antibiotics (day-3). After this initial transfer to Zn amended media, cells were passaged once a day into fresh Zn amended media via 1:100 dilution in until day zero. At day zero, both groups were tested for susceptibility to ciprofloxacin as well as their susceptibility to Zn via standard broth microdilution assays described above. As an exposure-based control, cells passaged for five days but not previously exposed to Zn were also tested. These wild type, zero days Zn pre-exposure cells would serve as a control for the impact of Zn pre-exposure on resistance development. A schematic of the experiment is displayed below in Figure 1A.

**Figure 1.**
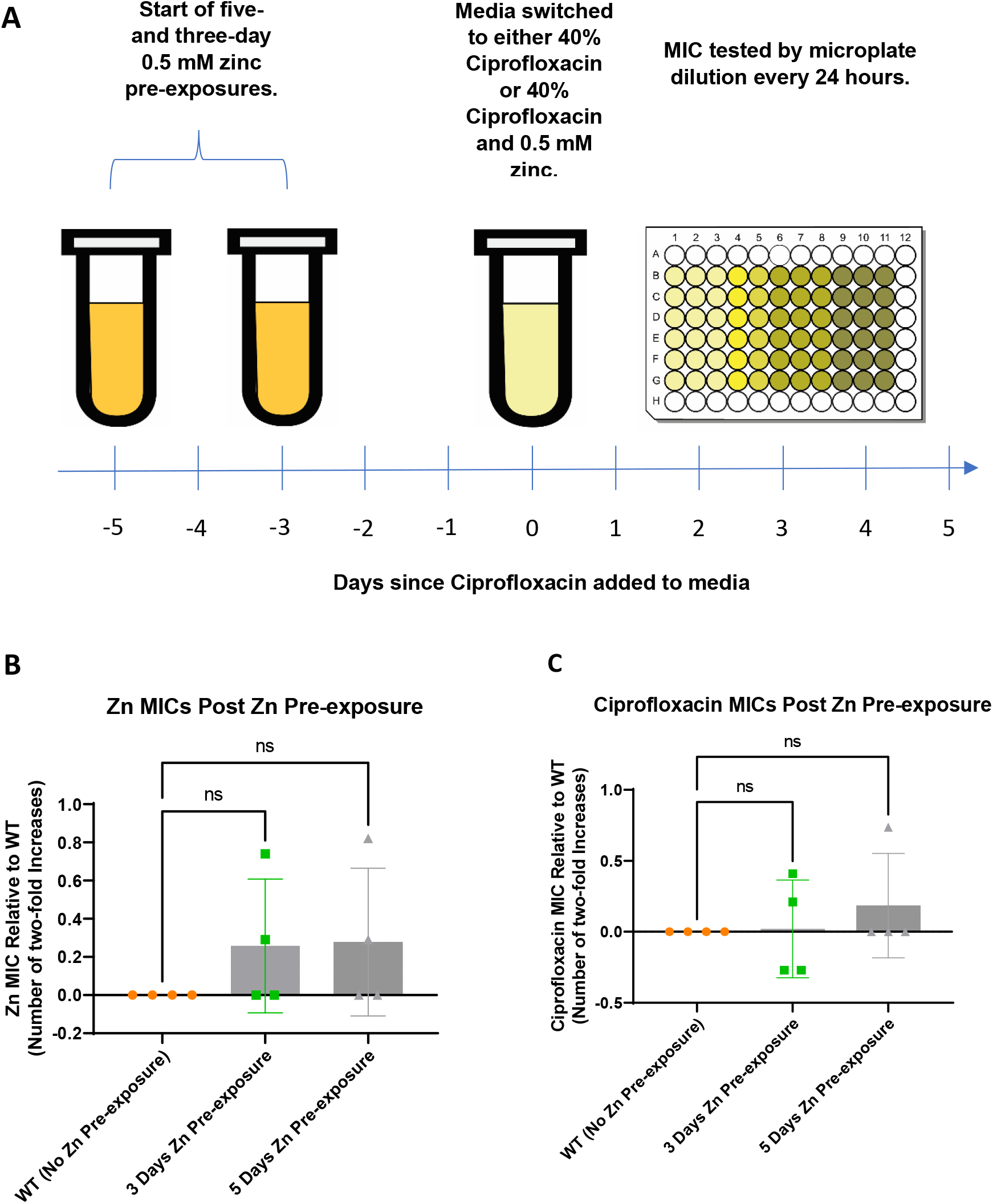
Fold Change in MIC after Zn only pre-exposure period. A). E. coli was pre-exposed to zinc for three or five days. On day zero, pre-exposed cells along with wild type, zero-day pre-exposed cells, were exposed to subinhibitory ciprofloxacin and tested daily for resistance development by assessing their MICs. E. coli exposed to Zn for up to five days showed no significant change in susceptibility compared to unexposed, control samples as assessed by MIC measurement for Zn (B), and ciprofloxacin (C). In both cases, results are plotted as averages of biological replicates with error bars representing the standard deviation of the samples with N=4.

In addition to the direct impact that Zn pre-exposure has on resistance development, tests were also performed to examine if the impacts persisted after the removal and substitution of Zn from the bacterial environment with subinhibitory concentrations of ciprofloxacin. This was tested by splitting the 5 days Zn exposure 3 days Zn exposure and 0 days Zn exposure control conditions 1:100 on day zero into two separate groups: 1) a Zn and antibiotic combination group containing 4mL of LB broth still supplemented with 0.5 mM of Zn as well as ciprofloxacin at 40% of its MIC concentration and 2) 4mL of LB broth supplemented with only ciprofloxacin at 40% of its MIC concentration. After 24 hours, 40 μl of cells from each exposure condition was added to 4 mL of fresh media containing the corresponding treatment (i.e., cells exposed to Zn for 5 days and then exposed to a combination of 0.5 mM Zn and 40% ciprofloxacin MIC were sub-cultured into fresh media containing 0.5 mM Zn and 40% ciprofloxacin MIC) for another 24 hours. This was repeated for seven days with antibiotic susceptibility tested after days 1, 2, 3, and 7. Experiments were performed independently in biological quadruplicate.

### Whole Genome Sequencing

Experimental samples to be sequenced had single colonies selected via 24-hour growth on agar plates. This was done for three biological replicates that were pre-exposed to Zn for five days with the purification occurring at the D0 timepoint. After selection, these colonies were transferred to 4mL of LB and grown to saturation and stored at −80C for future use. DNA extraction and whole genome sequencing of cells were performed by Microbial Genome Sequencing Center (Pittsburgh, PA) at a depth of 200Mbp on the Illumina NextSeq2000 platform. Data processing was performed using CLC Genomics Workbench (Qiagen). Fastq files containing sequencing reads from paired end reads, were aligned to the reference *E. coli* MG1655 genome FASTQ file (NC_ 000913) downloaded from NCBI. Detection of mutations (SNPs, insertions and deletions) was performed using CLC Genomics Workbench (Qiagen).

### Assays for mechanism of resistance development and stability of resistance

After sequencing and purifying the three samples which were pre-exposed to zinc for five days, these three D0 samples were either grown in LB then re-exposed to 40% ciprofloxacin and had their MICs assessed each day following the same protocols stated previously or were preexposed to Zn for five days prior to being exposed to 40% ciprofloxacin following the protocol described in the Zn pre-exposure section. This was done to assess if the observed mutations were responsible for the change in the dynamics of resistance acquisition. In both cases, MICs were compared against WT samples that were not previously exposed to Zn before their exposure to ciprofloxacin.

We lastly checked to see if the post ciprofloxacin elevated resistance levels observed in the pre-exposed Zn samples were stable. Previously frozen samples that displayed elevated MICs were inoculated in LB media and grown for 24 hours until saturated. These samples had their MICs reevaluated using the standard broth microdilution MIC method used throughout this manuscript which were compared against the original values observed.

### Persistence of Zn’s effects on MIC

Saturated liquid cultures were diluted 1:100 in 4 mL of LB broth. Cells were then diluted 1:100 into 4 mL of LB broth containing an additional 0.5 mM of Zn. Cells were kept in this Zn pre-exposure condition for five days with daily passaging of 40uL into 4mL of fresh media. After five days of Zn pre-exposure, cells were transferred to unamended LB media for one and three days before being exposed to LB amended with 40% ciprofloxacin. Each condition had its MIC assessed daily after exposure to ciprofloxacin. As a control, cells not previously exposed to Zn were also tested with no delay between Zn pre-exposure and ciprofloxacin exposure. MICs were tested each day.

### Growth metrics and tolerance

*E. coli* samples that were pre-exposed to Zn for five days had their growth examined. After preexposure concluded, these samples and unexposed, WT counterparts were diluted 1:100 in 100uL of LB media supplemented with 40% ciprofloxacin on a 96 well plate covered by its lid to mimic their first day of growth in a ciprofloxacin environment. To prevent condensation, plate lids were made hydrophilic by pouring 2 to 3 ml of 0.05% Triton X-100 in 20% ethanol into the cover. The surface was coated by tilting the lid several times to ensure even coverage of the inner surface. After 15 to 30 seconds, the treatment solution was poured off, and the cover was shaken to remove most of the remaining liquid before being allowed to air dry. Absorbance at OD600 was assessed overnight every five minutes for 16 hours in a plate reader at 37 C.

Readings were graphed in Microsoft Excel to determine both the doubling time and the lag time for each condition. To help ensure uniformity, we defined lag time as the length of time for the sample to reach an OD600 value of 0.01 after normalization to its starting inoculum reading. Statistical analysis was performed in GraphPad Prism 9.

*E. coli’s* tolerance to ciprofloxacin was also assessed after Zn pre-exposure by performing a time-kill assay^26^. Samples were serially passaged during five days of pre-exposure in either LB or LB supplemented with 0.5mM zinc. After the completion of the pre-exposure period, samples were diluted 1:100 into 4mL of LB containing a high concentration of ciprofloxacin (18 times the MIC) and incubated at 37C and 180rpm. The concentration of live cells was enumerated via plating on agar just prior to incubation (0 hours) as well as after 0.5, 1, 1.5, 2, 3, 4, 5, and 6 hours of incubation. The fraction of surviving cells for both condition was then plotted in GraphPad Prism on a semilog scale to examine both conditions’ killing rate.

## Results

### Prolonged Zn exposure alone does not impact ciprofloxacin or Zn resistance

The ciprofloxacin MIC for *E.coli* was determined to be between 0.0375 and 0.01875 ug/mL (upper bound 0.0375 ug/mL used) and the Zn MIC for *E.coli* was determined to be between 1.5 mM and 1mM (upper bound 1.5 mM used), both of which were in the expected MIC range from the literature^3,18^. We next examined if ciprofloxacin and Zn combined had any interactions by assaying the fractional inhibitory concentration (FIC) of the two together. The FIC was 1.5, indicating that ciprofloxacin and Zn did not have a synergistic nor antagonistic interaction but was indifferent. Thus, Zn was not expected to interfere with the activity of ciprofloxacin.

We first investigated the impact of continued Zn exposure on resistance, to mimic a multi-day course of Zn supplementation. We selected 0.5mM as the exposure concentration based on a prior study in the literature, as well as to maintain a similar % MIC to that of ciprofloxacin used in subsequent experiments^19^. Cells were exposed to 0.5mM of Zn for three or five days and the MIC to ciprofloxacin and Zn were assessed. We found that Zn pre-exposure did not significantly change the ciprofloxacin (Fig. 1B) or Zn MIC (Fig. 1C) relative to cells from a control passage in LB.

### Pre-exposure to Zn impacts ciprofloxacin resistance dynamics

While Zn exposure alone had no observable impact on ciprofloxacin resistance development, we next sought to determine whether subsequent resistance development in the presence of antibiotics was altered in cells that had been pre-exposed to Zn. This experimental design models treatment cases where a child is first given Zn/ORS treatments and then begins taking antibiotics, which often are acquired without need or recommendation from a doctor. Thus, cells that were pre-exposed to 0.5mM Zn for 5 or 3 days were subsequently exposed to ciprofloxacin at a concentration of 40% of its MIC. The MIC of ciprofloxacin was then measured over a week of exposure. Two post Zn pre-exposure groups were assessed: a single antibiotic group and a combined antibiotic and Zn group. In the single antibiotic group, after the Zn preexposure period ended, the cells were exposed to only ciprofloxacin and their media was no longer supplemented with Zn. This group represented the condition where Zn/ORS treatment was completed or otherwise stopped before the onset of antibiotic consumption. In the second combination group, after the Zn pre-exposure period was concluded, cells were exposed to ciprofloxacin and Zn combined, representing conditions where Zn/ORS was continued while antibiotics were consumed.

We observed that prolonged Zn pre-exposure led to faster increases in resistance (MIC) after exposure to ciprofloxacin alone, compared to no Zn-pre-exposure (Fig. 2A), with a significant difference observed on days two (Fig. 2B) and three (Fig. 2C) between the five day Zn preexposed condition and the zero days Zn pre-exposure condition via ordinary one way ANOVA with multiple comparisons (day two: p= 0.0049, day three: p =0.0197). However, after a week of exposure to ciprofloxacin these differences were no longer observed, with no statistical difference in MICs between the three groups. This suggests that Zn pre-exposure does not elevate the MIC beyond the level it would reach in the absence of Zn pr-exposure, but accelerates resistance development.

**Figure 2.**
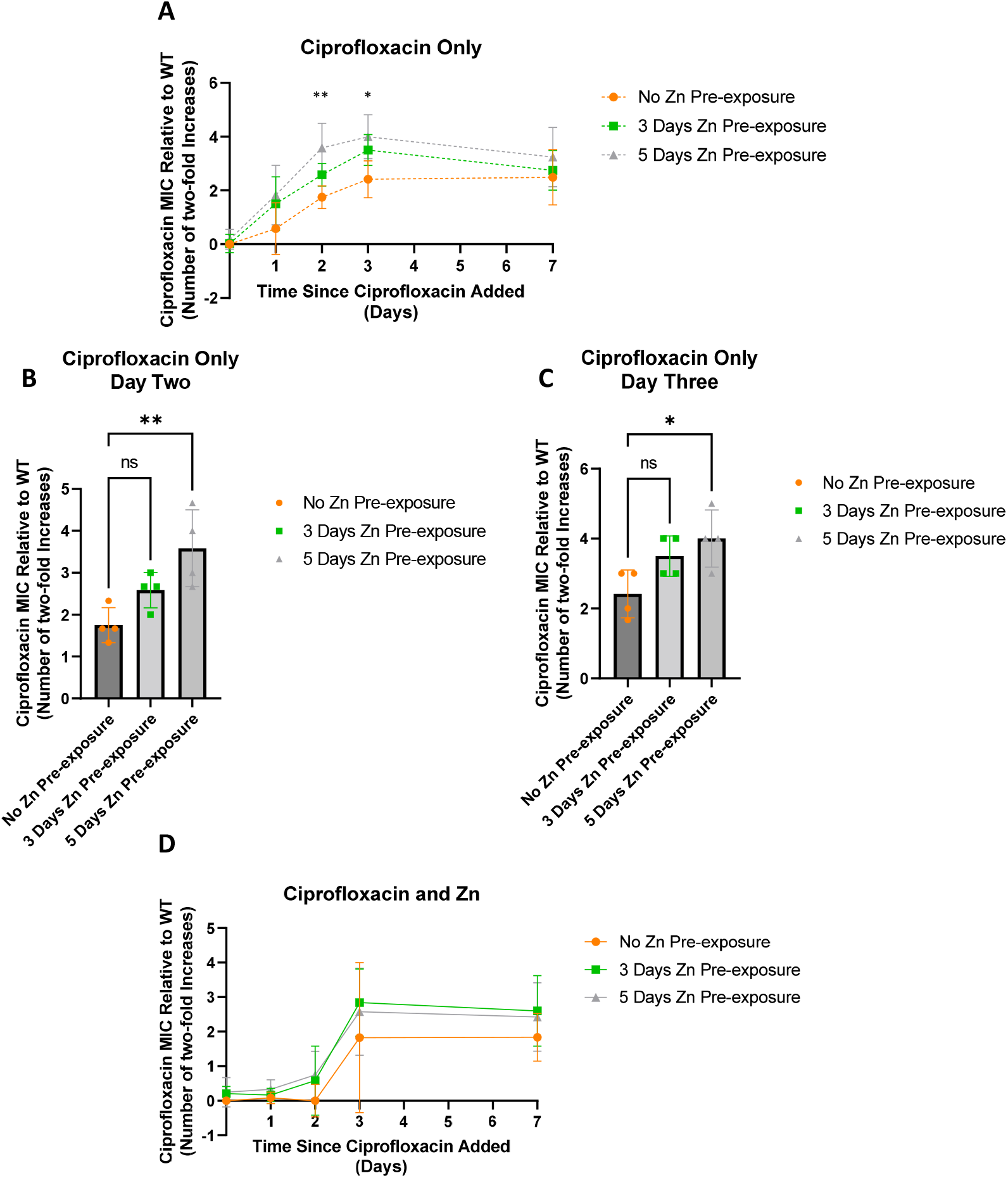
Change in ciprofloxacin MIC overtime. After three days of exposure to only subinhibitory concentrations of ciprofloxacin, E. coli pre-exposed to 0.5mM Zn for five days showed higher MICs that were significantly different compared to samples not pre-exposed to Zn two and three days after exposure to ciprofloxacin (A). While a significant difference was observed for the five-day Zn pre-exposure samples on days two (B) (p=0.0049) and three (C) (p=0.0197), no significant difference was ever observed for the three-day pre-exposure samples. Meanwhile, samples exposed to a combination of subinhibitory ciprofloxacin and 0.5mM Zn after the initial 0.5mM Zn pre-exposure period saw no significant differences between pre-exposed conditions and the non-pre-exposed condition (D). Values are plotted in GraphPad Prism as the average of N=4 biological replicates with error bars indicating standard deviations for each data point. All comparisons are nonsignificant unless otherwise noted.

To better ensure that this difference was due to the five-day pre-exposure in zinc and not the additional passages, we compared the MICs of *E. coli* passed for 5 days in LB to cells not pre-passed in LB (Supplemental 1). Here we saw passage number had no statistically significant difference on the baseline (D0) MIC nor the MICs recorded after 1,2,3, and 7 days of ciprofloxacin exposure.

Unlike in the ciprofloxacin only condition, when Zn exposure was maintained beyond the original pre-exposure timeframe, no statistically significant differences were observed via one way ANOVA with multiple comparisons between the five- or three-day pre-exposure condition for any of the time points examined (Figure 2D).

### Ciprofloxacin and Zn together slow resistance development

After noting the different behaviors between the ciprofloxacin only group and combination exposure group for the pre-exposure time tests, we probed to see if there was any difference in resistance development between ciprofloxacin and zinc compared to ciprofloxacin only. Furthermore, we also texted if this difference changed for the different Zn pre-exposure periods (none, three and five days). We found that samples treated with Zn and ciprofloxacin together led to slower resistance development compared against ciprofloxacin only samples when pre-exposure time to Zn was controlled (Figure 3). Here, the presence of Zn in the Zn and ciprofloxacin conditions resulted in significantly lower relative MICs on day two when there was no zinc pre exposure (Fig. 3A: p=0.0015), three days of zinc pre-exposure (Fig. 3B: p=0.0101) and five days of zinc pre-exposure (Fig. 3C: p=0.0026). Furthermore, Zn pre-exposure did appear to impact this comparison as significant difference observed a day early on day one only for the pre-exposed samples (p=0.0394, p=0.0390 for Fig. 3B, Fig. 3C respectively). These results were largely expected, especially for the no Zn pre-exposure condition, as previous literature reported chelating interactions between ciprofloxacin and Zn that decrease cellular permeability to fluoroquinolones^27^. Furthermore, it was shown that Zn increased concentrations of ciprofloxacin required to select for resistance^19^. As was observed previously, after a week of exposure to ciprofloxacin all conditions became more uniform with no significant difference in MICs.

**Figure 3.**
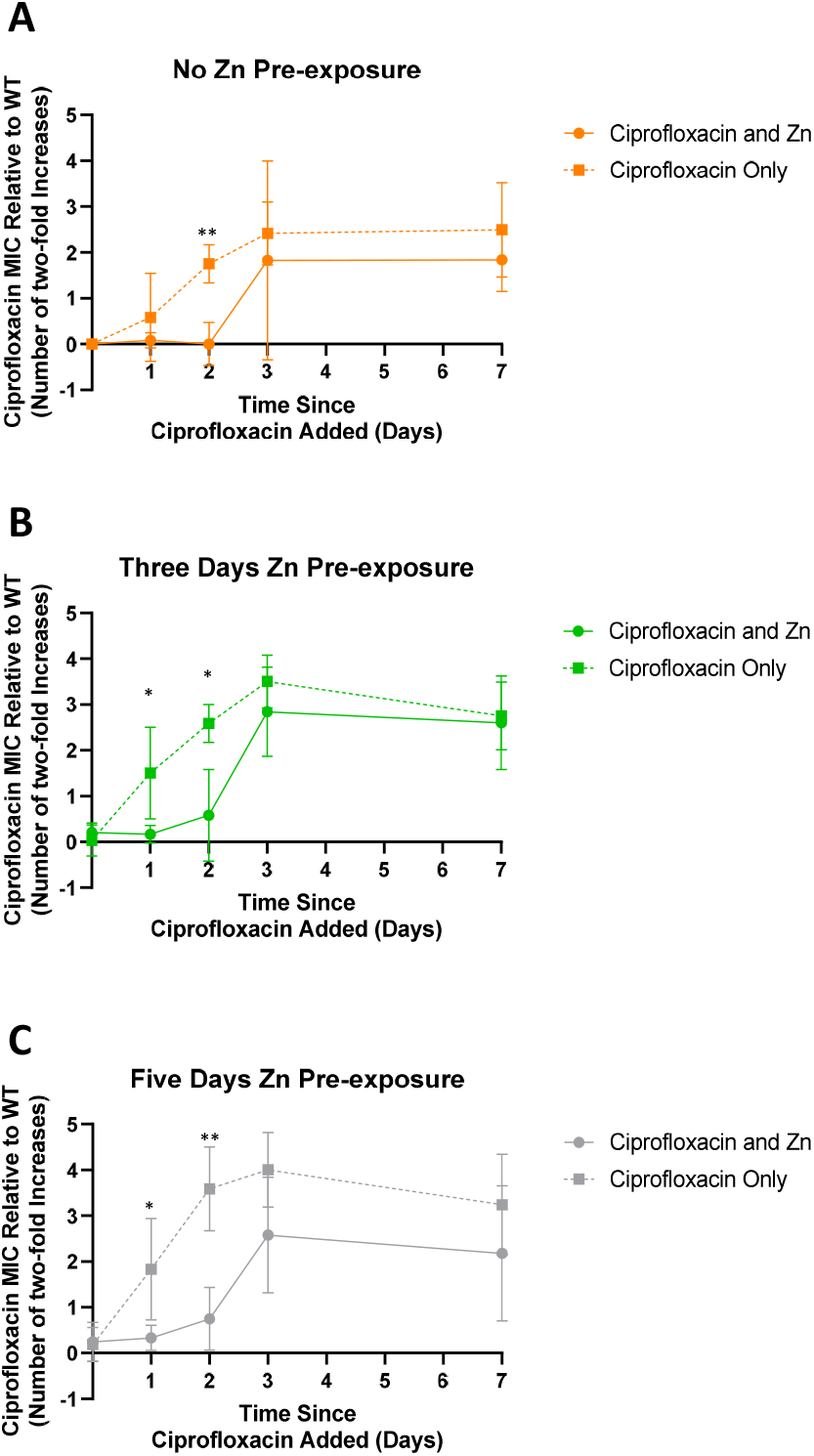
Comparison between Zn and ciprofloxacin combination and ciprofloxacin only exposure with Zn pre-exposure time controlled. The continued presence of Zn when E. coli is exposed to ciprofloxacin in the combination group leads to a slower increase in MICs for up to three days after the antibiotic is first introduced. This was observed to be statistically significant via unpaired t test when there was no Zn pre-exposure (A) on day two (p =0.0015), when there was three days of Zn pre-exposure (B) on days one (p=0.0394) and two (p=0.0101) and when there was five days of Zn pre-exposure (C) on days one (p=0.0390) and two (p=0.0026). Values are plotted in GraphPad Prism as the average of N=4 biological replicates with error bars indicating standard deviations. Unless otherwise noted, all other comparisons were nonsignificant.

### Zn pre-exposure resistance acceleration not due to underlying genetic changes

After observing these differences, we performed whole genome sequencing to discern any potential genetic changes relative to the previously sequenced WT that afforded cells the ability to gain resistance more quickly after Zn pre-exposure^3^. To do so, samples were purified after five days of Zn exposure but prior to ciprofloxacin exposure (day zero) to a single clonal population which was sent for sequencing. After discounting any mutations that fell below 90% frequency, the unique mutations displayed in Table 1 remained.

**Table 1.** Polymorphisms among sequenced samples. All mutations across the sequenced samples with over a 90% variant frequency.

| Mutations Unique to Zinc Pre-exposed *E. coli* MG1655 Genome [U00096] |  |  |  |  |  |  |  |  |  |
| --- | --- | --- | --- | --- | --- | --- | --- | --- | --- |
| Condition and Name | Position | Location Relative to Gene | Type | Ref. | Allele | Amino Acid Change | Count | Coverage | Frequency |
| Sample #1<br>(Fim Region Mutation Zinc Five Days Pre-exposure) | 4542720 | Noncoding region (Downstream fimE, upstream fimA) | Deletion | T | - | N/A | 40 | 42 | 95.24 |
| Sample #2<br>(No Mutation Zinc Five Days Pre-exposure) | No unique mutations observed |  |  |  |  |  |  |  |  |
| Sample #3<br>(Second No Mutation Zinc Five Days Pre-exposure) | 2346054 | CDS position 1190 within nrdA | SNV | A | G | E->G | 106 | 106 | 100 |

The following unique mutations were found: a deletion outside of any coding region between the fimA and fimE gene in sample #1 and a single nucleotide variation resulting in a change from a glutamic acid to a glycine in the coding region of the nrdA gene in sample #3. With the genetic changes among these samples known, these three previously Zn pre-exposed samples were exposed to sub-inhibitory ciprofloxacin, to see if any of the mutations were responsible for the accelerated elevated resistance phenotype. We found that all samples behaved similarly to each other regardless of genetic composition and none showed accelerated resistance previously observed for cells pre exposed to five days of Zn (Figure 4). Notably, these samples were subject to outgrowth in the absence of Zn before re-testing. Thus, accelerated resistance previously observed (Fig. 2) does not appear to be due to an altered genotype. Instead, it may be due to expression or metabolic changes occurring immediately after Zn pre-exposure.

**Figure 4.**
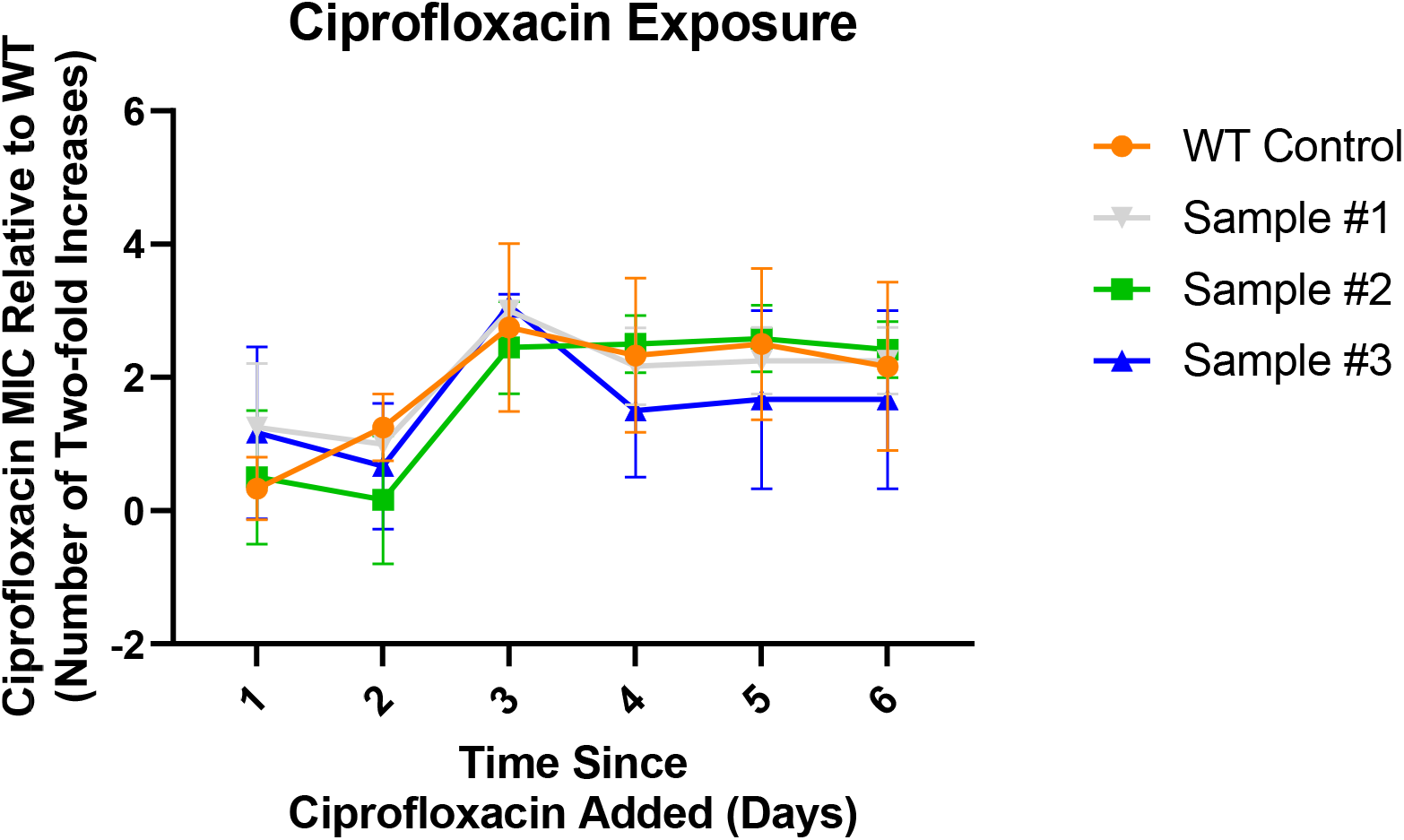
Impact of mutations from prior Zn exposure on ciprofloxacin development. All samples previously pre-exposed to Zn before being frozen, showed no significant difference in the development of resistance to ciprofloxacin after being exposed to subinhibitory ciprofloxacin after being thawed. An ordinary ANOVA with multiple comparisons was performed between each sample and a control sample that was never exposed to Zn. Data represents the average MIC where N=4 with error bars representing the SD.

To determine if the samples needed to be directly exposed to Zn first to observe accelerated resistance development, the sequenced isolates were again subjected to the original Zn pre-exposure and subsequent ciprofloxacin exposure regimen. As an additional control, a previously wild-type condition was also exposed to zinc. This single Zn exposure control was included to probe if any of the sequenced samples behaved differently with the “extra” prior Zn preexposure, or specific genetic change. These samples were all compared against a similar no Zn pre-exposure, negative control as before.

As shown in Figure 5, all Zn exposure samples developed elevated MICs within the first two days before stabilizing at values similar to the WT control. All samples previously exposed showed larger relative MICs. These larger values were statistically significant relative to the MIC of the WT control on day one (Fig. 5B). Though the single Zn exposure sample had a higher MIC on day one, the difference was not significant with a p-value of 0.054. By day two (Fig. 5C), a significantly elevated MIC for the single Zn exposure relative to the WT control was observed, consistent with our five-day Zn preexposure result presented earlier (Fig. 2C). The accelerated resistance upon re-exposure to Zn not observed in Figure 4 suggest that the immediate role that Zn’s pre-exposure impact plays in the development of resistance development is more nuanced and not mutation-based, as the same samples only displayed the previously observed elevated behavior when the *E. coli* was again exposed to Zn directly prior to antibiotics. Overall, this seems to indicate that while Zn’s effects can be reversed, immediate usage of antibiotics after a course of Zn supplementation that did not help clear their diarrhea symptoms can negatively impact recovery, though further studies are warranted to expand this conclusion to *in vivo* environments.

**Figure 5.**
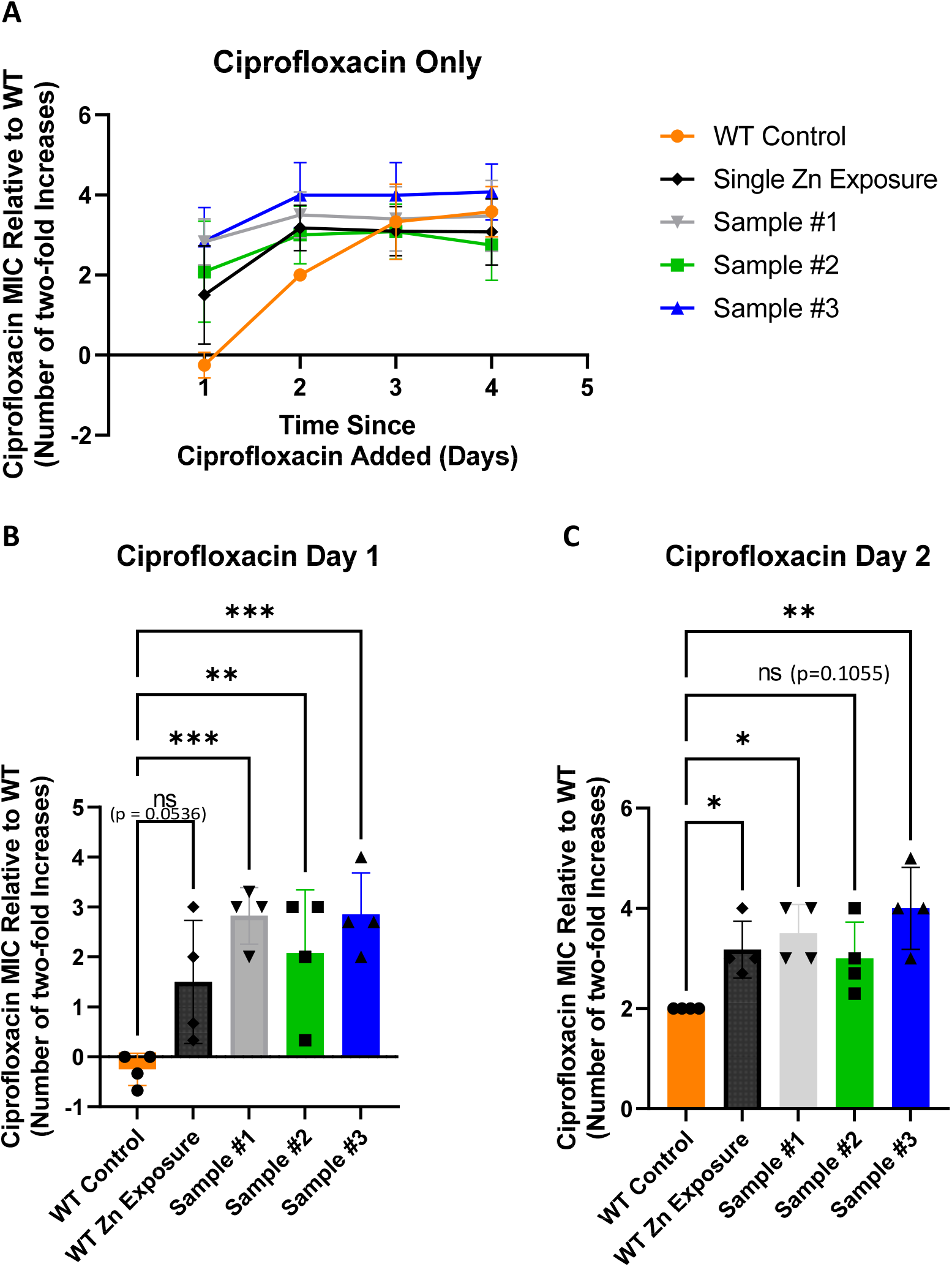
Re-exposure to Zn after freezing. To further examine if the earlier development of resistance among the Zn preexposed samples is a result of the specific mutations observed post Zn exposure, the same three samples previously exposed and sequenced along with a fresh wild type sample were thawed and pre-exposed to Zn for five days before being exposed to ciprofloxacin at a concentration of 40% of the MIC. Pre-exposing the samples to Zn post freezing was the only experimental difference between this figure and Figure 6. This was done to directly recreate the original experimental conditions. Here five days of Zn pre-exposure led to an earlier onset of elevated MIC levels (A). After one day of exposure to ciprofloxacin (B) all prior pre-exposed conditions saw significantly higher MICs compared to the control (Sample #3 p=0.0009, Sample #2 p = 0.0092, Sample #1 p = 0.0010). The newly exposed Zn exposure condition saw an elevated MIC but was not significant (p=0.0536). By day two (C) this newly Zn pre-exposed sample was significantly higher (p=0.0495) with differences still observed for Sample #3 (p = 0.0011) and Sample #1 (p = 0.0112). Although Sample #2 was still elevated, it was no longer significantly elevated compared to the control (p= 0.1055).

#### Accelerated elevated resistance is stable

After observing that Zn’s ability to cause resistance to develop earlier is non-permanent, we decided to examine if the faster enhanced resistance level displayed a similar degree of transience or was more stable. To do so we passaged samples from our Day 2 results from the prior experiment (Fig. 5C) (after all had previously shown elevated resistance levels compared against the control) in drug free media for one day and checked the MIC values. This timespan was equivalent to the amount of time it took for the zinc pre-exposure phenotype to no longer be observed. These retested MICs were compared against the original values recorded. As shown in Figure 6 below, in all four cases the re-examined MIC showed no statistically significant differences when compared to the original value. This lack of change in relative MIC indicated that the resistance that did develop was stable. This furthers motivation for similar studies to be performed *in vivo* to see if the gut environment of a child taking Zn, followed by antibiotics recreates these conditions as there is concern that if these conditions were recreated and observed in vivo, these patients infections might become resistant before they are able to be cleared by current treatment approaches and the bacteria would keep their resistant phenotype. This could serve as a reservoir for antibiotic genes to be passed to other future infections within the host while also leading to shedding of these resistant bacteria into the environment. This could be especially problematic in conflict settings where sanitation systems are lacking and might fuel more severe infections in the future. Similar *in vivo* findings would ultimately serve as an indication that current Zn/ ciprofloxacin practices should be reviewed as the impact of Zn would be more nuanced and not merely a silver bullet for diarrhea treatment.

**Figure 6.**
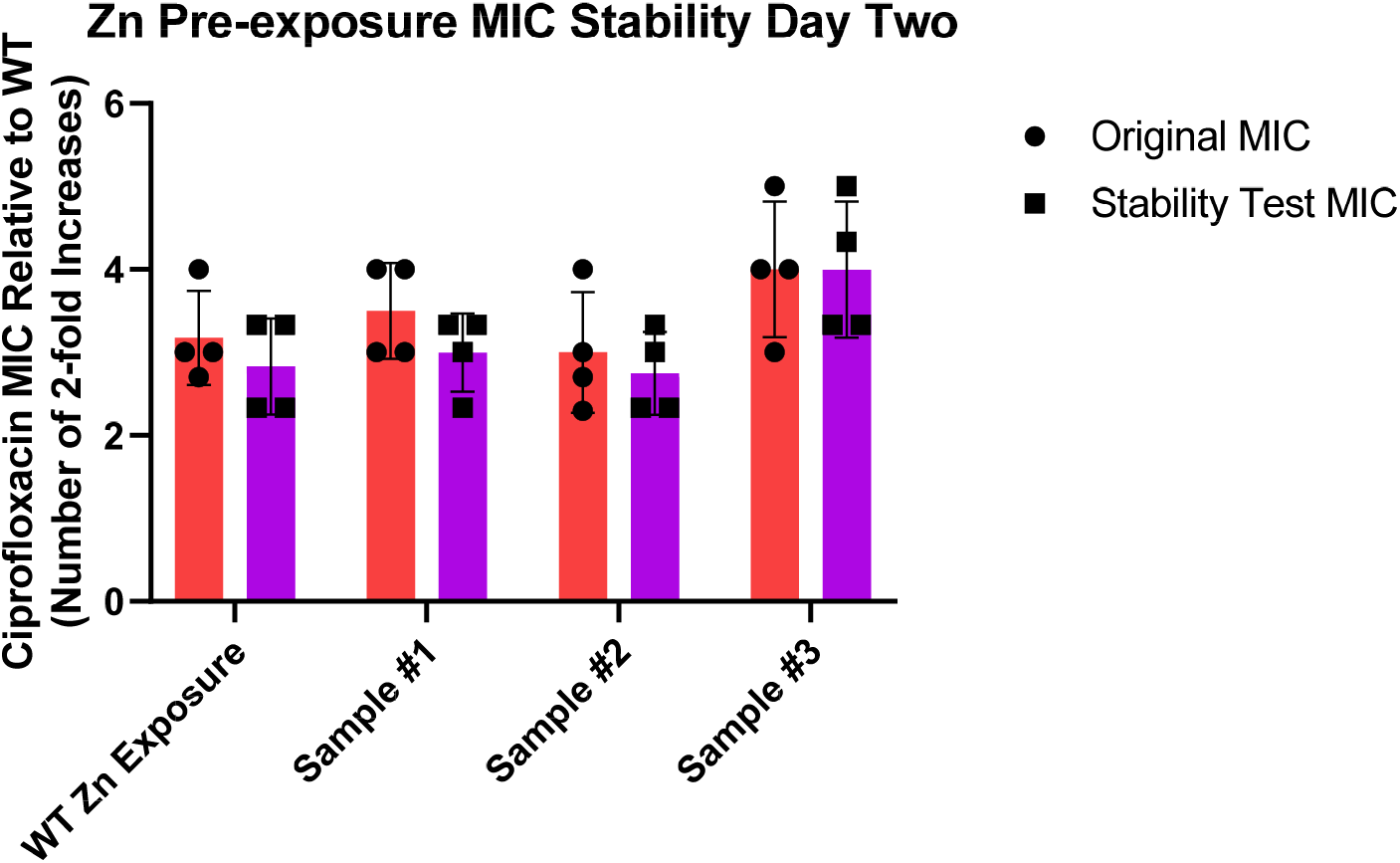
Resistance stability tests for samples pre-exposed to Zn. Comparisons between the original MIC observed and MICs tested three weeks later for the wild-type Zn exposure sample, Sample #1, Sample #2, Sample #3. In all cases no significant difference was observed between the samples which are graphed in GraphPad prism as the mean of four biological replicates with error bars representing the standard deviation.

### Impact on Growth and Death Dynamics

To further probe a possible underlying mechanism, the growth curve of the Zn pre-exposed samples was determined. Growth over a 16-hour time frame was examined to see if the elevated MICs resulted from cells doubling faster in ciprofloxacin conditions and thus ultimately allowing for comparatively more mutations to occur due to the existence of more cells in the ciprofloxacin environment. Lag time was assessed for a similar reason and to see if these pre-exposed cells were able to adapt to their environment more quickly. Results in Figure 7 indicate that within the 40% ciprofloxacin media, samples that were pre-exposed to Zn for five days ultimately had a significantly longer doubling time and lag phase compared to cells not pre exposed to zinc. After observing a slower growth rate and an increase lag, which could be indicative of the emergence of tolerance as reported by literature, we performed a kill curve to see if samples were inherently more tolerant to ciprofloxacin^28,29^. Despite the previously observed increase in lag time, samples show no statistically significant difference at viable colony counts at each time point measured, indicating similar kill rates from ciprofloxacin between the control and Zn pre-exposed conditions up to six hours after exposure.

**Figure 7.**
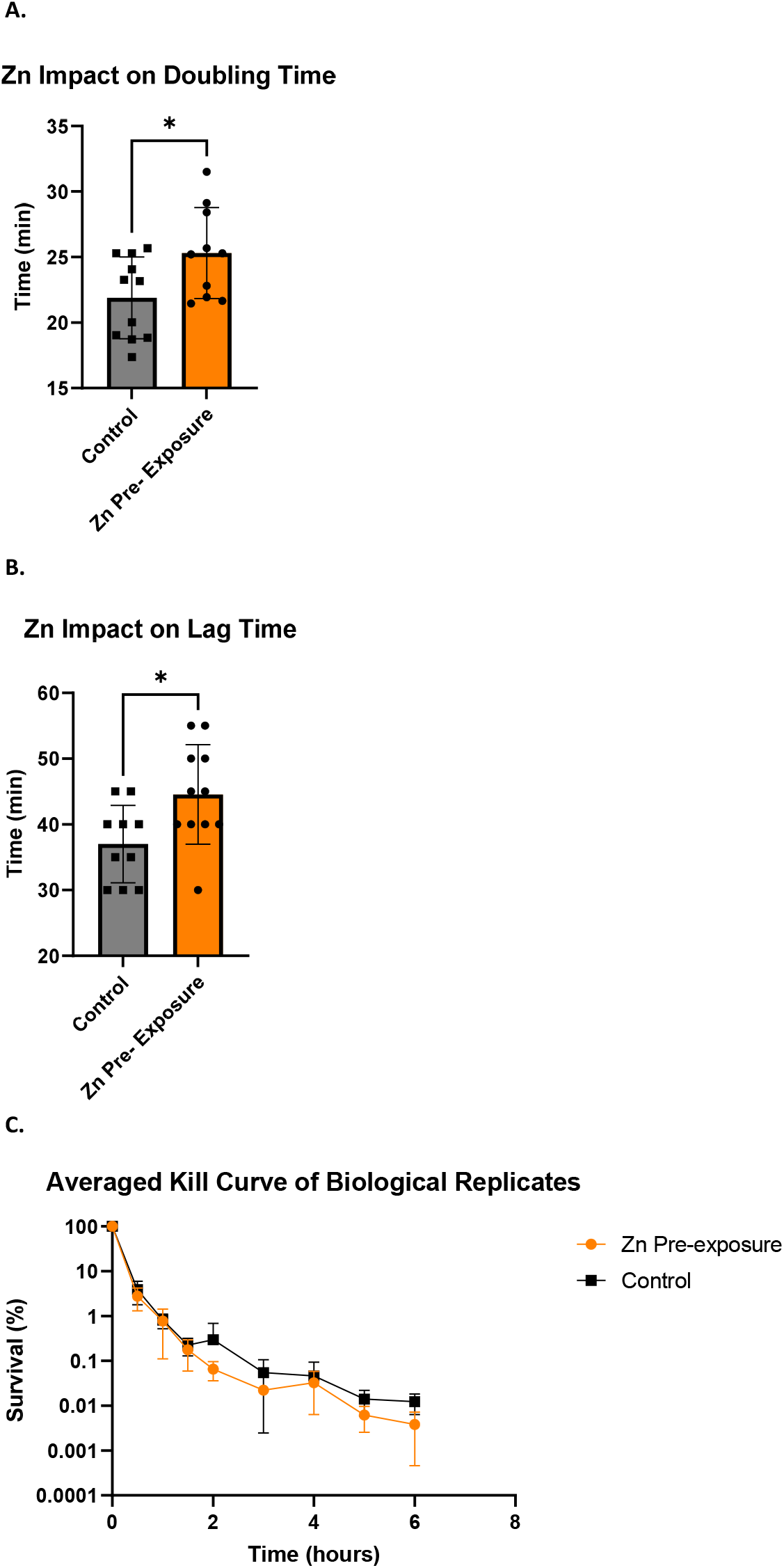
Comparison of Growth metrics for Zn pre-exposed samples. Five-day pre-exposed samples from stationary phase were diluted 1:100 in 40% ciprofloxacin with OD600 readings taken every five minutes. A) The doubling time of E. coli was measured and compared under both conditions where it was observed that samples pre-exposed to Zn grew slower than their WT counterparts. B) Likewise, these pre-exposed cells also displayed a longer lag phase which was measured as the amount of time for the OD600 value to reach 0.01 when normalized to its starting value. C) No difference was observed in survivability after six hours of exposure to high concentrations (18x MIC) of ciprofloxacin.

Overall, our study shows that the timing and order of Zn pre-exposure relative to ciprofloxacin exposure plays a role in *E. coli’s* resistance development. Specifically, we observed that five days of Zn pre-exposure followed by subinhibitory ciprofloxacin led to elevated MICs faster than samples exposed to only subinhibitory ciprofloxacin. While the exact underlying mechanism has not yet been elucidated early results indicate the zinc does not result in a genetic change nor enhanced tolerance to ciprofloxacin, although it does result in slower growth and a longer lag phase in the subinhibitory ciprofloxacin environment compared to samples not pre-exposed.

### Discussion and Conclusions

Our study shows that Zn has a “multidirectional” impact on the development of resistance to ciprofloxacin. More specifically, we note that extended pre-exposure to Zn alone followed by ciprofloxacin accelerates development of resistance early on, but co-exposure to ciprofloxacin and Zn delays resistance development. While the complete mechanism is not yet known, the lowered MIC observed for combinations of Zn and ciprofloxacin could be due to a few factors such as the addition of zinc inhibiting the SOS response and associated hypermutation abilities in E. coli^23^ or even Zn chelating the ciprofloxacin in the media when together, altering the effective concentration of ciprofloxacin which prior studies have shown can help select against resistance^19,27^.We note that it is also possible the binding could occur within the *E.coli* itself given prior studies on modeling Zn transport have noted the existence of a reservoir of unbound Zn within the cell cytoplasm^30^. While this split in behavior is interesting and merits further future studies, it is beyond the scope of this paper. In this study, we focus on the observed negative impact that zinc pre-exposure has on resistance development and attempted to provide insight into this phenomenon. While these elevated MICs were observed to not be due to a genetic mutation, they could be the result of an epigenetic change that persists for a number of generations even after the stressor (in this case the Zn) is removed from the environment such as the aggregation of misfolded proteins which have been shown to form in *E.coli* as a result of proteotoxic stress and provide protection from additional stressors including antibiotics such as streptomycin^31^. Alternatively, this observed behavior could be due to an alteration in expression level of transport genes associated with resistance development such as the *mdt* operon encoding a multi-drug resistance pump, or enhanced expression of zinc transporters, both of which have been shown to be over expressed after a single overnight exposure to zinc sulphate^32^. The mechanism underlying the altered resistance dynamics is an area for future studies as our results show that it is not due to genetic changes and there is no change in tolerance to ciprofloxacin despite an increase in lag time (and decrease in growth rate).

While the specific mechanism and generality of this phenomenon warrant additional *in vitro* studies, we believe that this study provides clear evidence of the importance of examining if these observations are repeatable during *in vivo* studies. Although we focus on zinc supplements as part of a disease treatment prior to the use of antibiotics, given the high prevalence of oral supplement usage around the world, similar interactions are possible. Understanding these mechanisms could have important outcomes for modified usage and prescriptions of supplements and antibiotics examining not just the interactions that exist between supplement and medication when taken together but also any “lingering” impacts that might result from a history of taking a specific supplement before any medication. While this work provides some evidence that despite Zn’s large acceptance for use in cases of diarrhea, more nuanced studies might need to be performed to see if these Zn treatments are promoting AMR development or enhancing the speed with which it develops in the gut. This is especially true for diarrhea cases that become persistent which would result in the recommendation for the use of antibiotics after the zinc treatment has already been deployed. This could help to provide a more complete picture of the role of Zn in the treatment of diarrhea diseases and serve to ensure that these children who suffer from such a major source of mortality are not having their condition inadvertently made worse by potentially outdated medical advice.

## Acknowledgements

We appreciate discussion of various aspects of the project with Professor Antoine Abou Fayad (American University of Beirut), Dr. Najwa Dheeb (UNICEF Yemen) and Professor David Hamer (Boston University). Funding for this project came from discretionary research support to Professor Zaman from Boston University.

